# ClearX9^™^: an efficient alternative to fetal bovine serum for growing animal cells in vitro

**DOI:** 10.1101/2023.05.01.538513

**Authors:** Sumit Gautam, Neeraj Verma, Siddharth Manvati, Pawan K. Dhar

## Abstract

Fetal Bovine Serum (FBS) is a nutrient-rich fluid that contains nutritional and macromolecular factors essential for cell growth. Every year millions of pregnant cows are slaughtered in search of FBS leading to huge environmental consequences. Here we report ClearX9™ - an affordable, sustainable, ethical, and effective replacement for FBS. ClearX9™ cell culture medium was used to grow HeLa (cervical cancer cells), HEK293T (embryonic kidney transformed cells) and Nthy Ori-3-1 (primary thyroid follicular transformed epithelial cells) and showed encouraging growth patterns and good cellular health. Compared with the FBS-enriched cell culture medium, ClearX9™ scored positive on all the parameters suggesting ClearX9™ as a credible alternative to FBS. In future, more work is required to establish the efficacy of ClearX9™ in toxicology testing, bio-manufacturing, regenerative medicine, and vaccine research.

**HIGHLIGHTS:**

- ClearX9™ provides good nutritional support for the growth of animal cells
- ClearX9™ cell growth performance is comparable to the serum-enriched medium
- ClearX9™ maintains a healthy morphological profile of cells during division
- ClearX9™ generates a stress-free environment within cells
- ClearX9™ does not require animal slaughter and reduces carbon footprint
- ClearX9™ has applications in biotechnology and cell cultivated meat industry

## 1. INTRODUCTION

Fetal Bovine Serum (FBS) is a standard supplement used in the animal cell culture medium providing a range of nutrients, growth factors, and other biological components for supporting the growth, proliferation, and differentiation of cells.

FBS is used extensively as a culture media supplement to culture cells from various animal and human tissues, including cancer cells, stem cells, and primary cells. FBS is used to culture cells that are genetically engineered to produce monoclonal antibodies, recombinant proteins, and vaccines. FBS has also been used in vaccine manufacturing, toxicology testing, and cell therapy experiments.

The use of FBS was first reported in the scientific literature in 1955 when RJ Eagle and colleagues compared the growth of several types of cells supplemented with serum from different animals and found FBS better than other sera in supporting the growth of most cell types (Eagle, 1955).

Due to its ease of extraction, storage, and widespread availability FBS has gained widespread usage in animal cell culture labs and industries worldwide.

Though FBS is the most essential component in the culture medium (Castells-Sala et al., 2017; Subbiahanadar Chelladurai et al., 2021) FBS comes with several limitations: batch-to-batch variability, risk of transmitting pathogens to humans, and slaughter of a pregnant cow leading to immense suffering to the animal.

On average each bovine fetus yields approximately 500ml of FBS It has been estimated that more than two million bovine fetuses are used worldwide to produce approximately 800,000 L of FBS (Lee et al., 2022). This is over and above the environmental cost of grass-feeding, ruminating, and methane-producing cow breeding.

These and many more concerns have accelerated the search for a credible alternative to FBS and built a new cell culture standard for the sustained growth of animal cells. To address pressing environmental, ethical, and safety concerns, ClearX9™ was designed as a natural growth media to provide nutrients for stress-free cell growth in-vitro.

## 2. MATERIAL AND METHODS

### 2.1 Cell line and reagents

HeLa, HEK293T, and Nthy Ori-3-1 cell lines were obtained from American Type Culture Collection (Manassas, VA) and grown in DMEM and RPMI 1640 medium (Invitrogen) supplemented with 10% heat-inactivated fetal bovine serum and 1% penicillin-streptomycin (Invitrogen) at 5% CO2, 37°C under a humidified chamber.

ClearX9™ was designed to offer nutrients and growth factors under natural and non-slaughter conditions (Indian Patent No: 403271, date 14.12.2020). The MTT [3-(4,5-Dimethylthiazol-2-Yl)-2,5-Diphenyl-tetrazolium Bromide] dye (M2128) was obtained from Sigma Aldrich. Triple distilled water (Millipore) was used for preparing solutions.

All reagents used in the experiments were of the highest purity grade.

### 2.2 MTT Assay

The effect of standard 10% Fetal Bovine Serum (FBS) supplemented cell culture medium and the ClearX9™ was assayed on the proliferation rate of HeLa, HEK293T, and Nthy Ori-3-1 cells was determined using MTT assay. Briefly, all cells (5× 10^4^/ml) were plated (∼10,000 / well) in triplicate in 96 well plates after the minimum three passages of each cell line. The cells were incubated in the presence of 10% FBS with traditional cell culture medium or ClearX9™ in a final volume of 200 μl for 72 hours at 37°C in a 5% CO_2_ humidified chamber. Cells grown with 10% FBS containing traditional cell culture medium served as a control. At the end of the cell culture, 20 μl of MTT solution (5 mg/ml in 1X PBS) was added to each well and incubated for additional 5 hours. The MTT containing medium was discarded. The resultant formazan crystals were dissolved in 200μl of DMSO. The absorbance value (A) was measured at 570 nm by a microplate ELISA reader.

### 2.3 Cell morphology study

Cells (7× 10^5^/ml) were plated in 2 ml traditional complete medium (DMEM/ RPMI 1640 + 10%FBS) in 35 mm^2^ culture dishes served as a control or ClearX9™ Complete Medium for 24, 48, 72 hours at 5% CO_2_, 37°C under a humidified chamber after the minimum three passages of each cell lines. Bright-field microscopy was used for studying morphology in response to the presence of containing traditional cell culture medium or ClearX9™. All images were captured at 10X and 20X magnification using a bright field microscope (Zeiss Primovert inverted microscope).

### 2.4 Statistical analysis

Statistical analyses were carried out using Graph Pad Prism software. Experimental data were expressed as means ± S.D. and the significance of differences was analyzed by a one-way ANOVA test. Tukey’s test was used to determine the significance of observed differences whereas the Dunnett test for used for testing experimental groups against the control group concerning cell viability.

## 3. RESULTS

Cell viability was evaluated on HeLa (cervical cancer) cell line using an MTT assay. The cells were incubated in the presence of 10% FBS containing traditional (DMEM) cell culture medium or “ClearX9™-version’s” for 72 hours and grown at 37°C in a 5% CO_2_ humidified chamber. Cells grown with 10% FBS containing traditional (DMEM) cell culture medium served as a control in this experiment (Fig.1)

**Fig 1:**
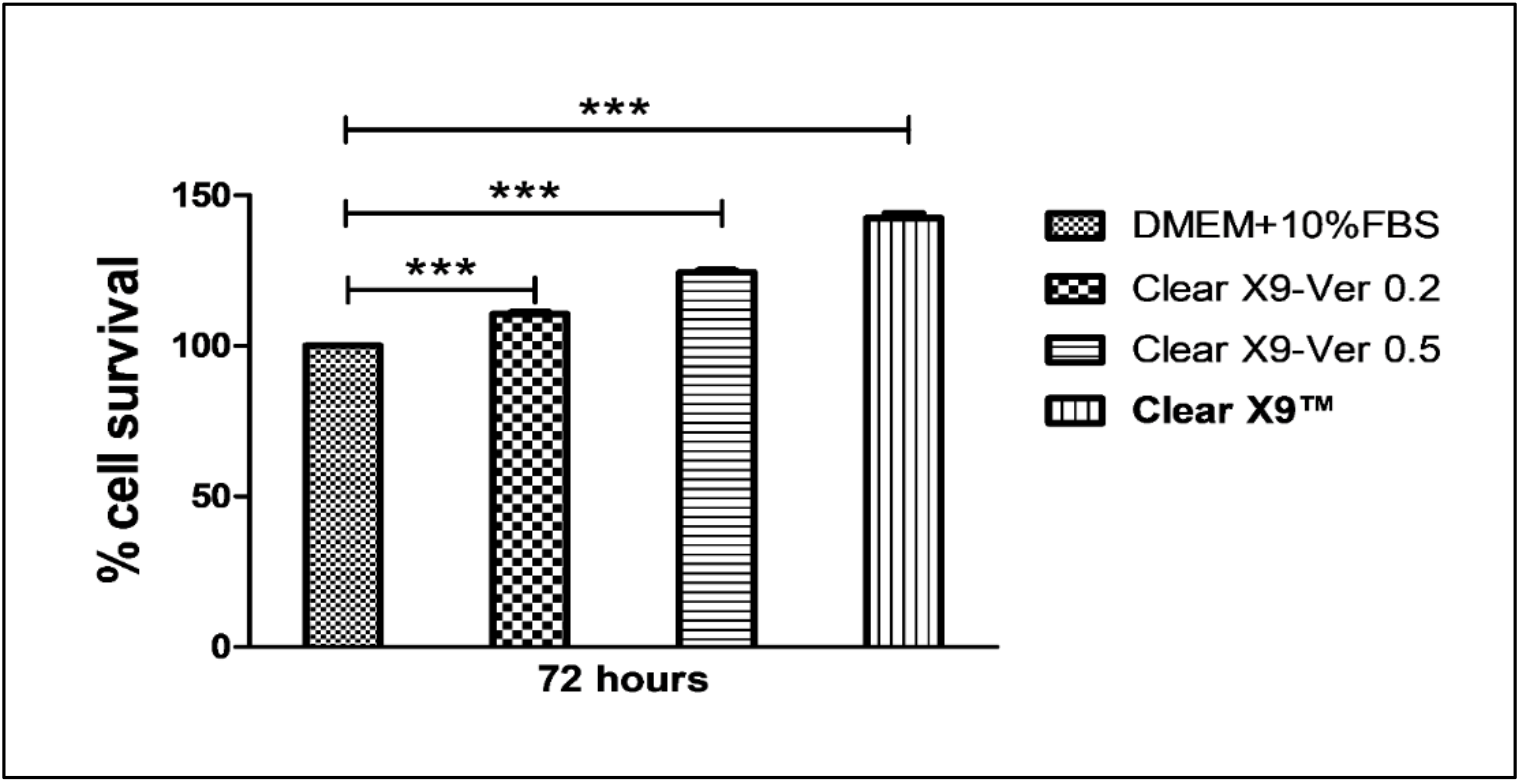
ClearX9™ mediated cells growth rate at 72 hours. The cell viability assay was also performed on HeLa cells using MTT assay. Briefly, cells (5 ×10^4^ cells/ml) were plated in 96 well plates and grown in the presence of ClearX9-Ver 0.2, ClearX9-Ver 0.5, ClearX9™ and Traditional Complete Medium (DMEM+10%FBS) for 72 hours. At the time of harvesting, MTT dye was added, and cells were incubated for 5 hours in a 37°C and 5% CO_2_ humidified chamber. The crystals formed were dissolved in DMSO and OD was taken at 570nM. The data were expressed in the form of mean values from three parallel experiments. Each experiment was performed in triplicate. ***p<0.001.

Data suggests increase in the proliferation rate of HeLa cells using ClearX9™ version (ClearX9-Ver 0.2, ClearX9-Ver 0.5, and ClearX9™) generating healthy cells profiles in all the versions of ClearX9™ in comparison to the 10% FBS enriched traditional (DMEM) medium (Fig. 1).

HeLa cells were plated at 7 × 10^5^ cells/ml in a 35mm^2^ culture dish and grown in the presence of 10% FBS containing traditional (DMEM) and three ClearX9™ versions (ClearX9-Ver 0.2, ClearX9-Ver 0.5 and ClearX9™) at 37°C in a 5% CO_2_ humidified chamber. HeLa cell morphology was studied after 72 hours of incubation using a bright field microscope. Images was captured by Zeiss Primovert microscope at 10X (Fig. 2A) & 20X magnification (Fig. 2B).

**Figure 2.**
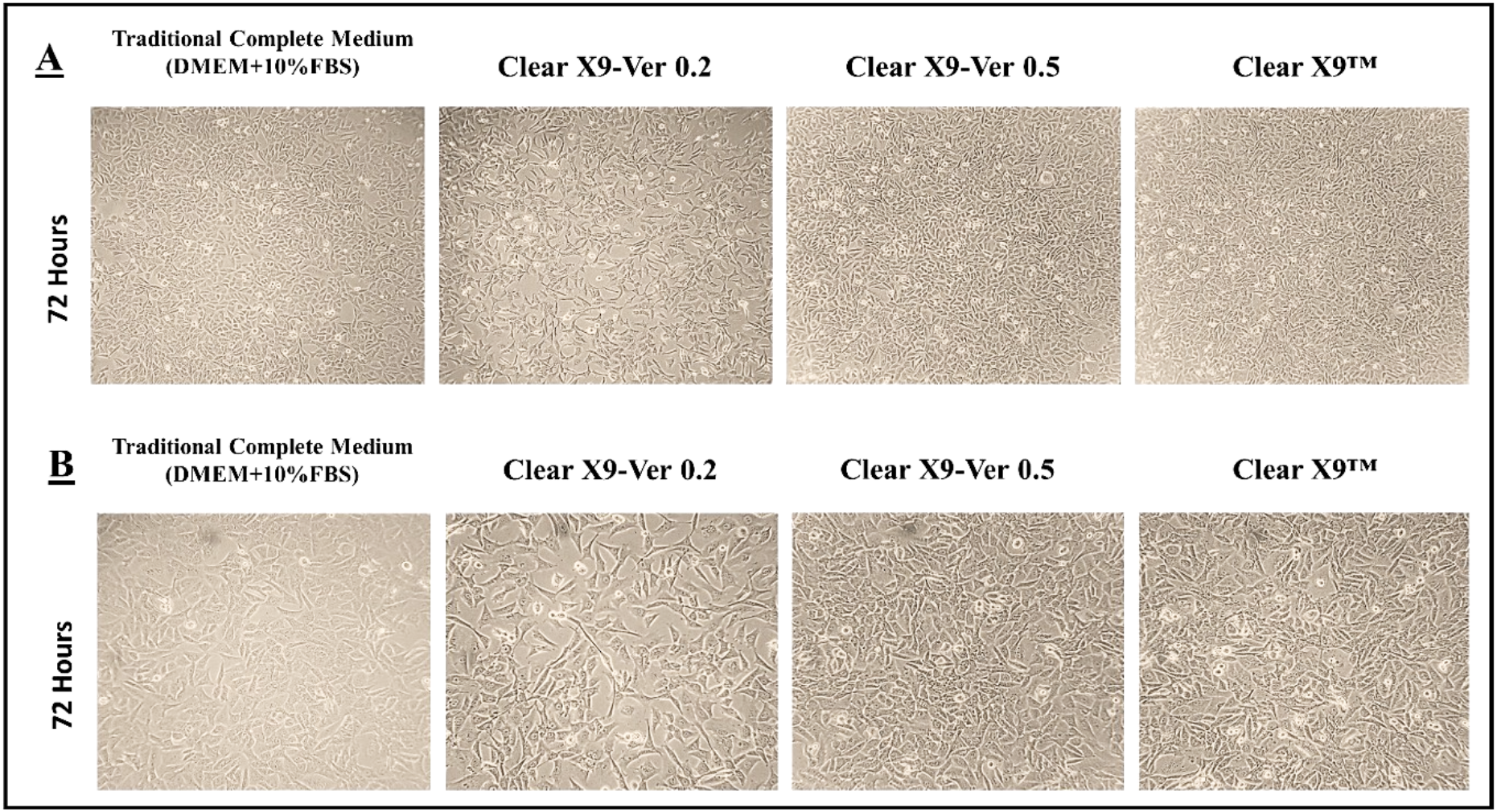
ClearX9™ induced growth rate of HeLa cells at 72 hours. 7 × 105 cells/ml cells were plated in a 35mm2 culture dish and grown in the presence of ClearX9-Ver 0.2, ClearX9-Ver 0.5 & ClearX9™ and Traditional Complete Medium (DMEM+10% FBS). The Cell Morphology was visualized after 72 hours using bright field microscope (Images were captured by Zeiss Primovert inverted microscope at 10X (Figure A) and 20X (Figure B) magnification).

ClearX9™ medium was found to increase the growth rate of HeLa cells in the presence of ClearX9-Ver 0.2, ClearX9-Ver 0.5, and ClearX9™ complete medium with cells showing healthy morphology compared to the 10% FBS enriched traditional (DMEM) medium cells (Fig. 2A, Fig. 2B). Compared to all the versions, maximal cell growth rate was observed in cells growing in ClearX9™. The cell viability pattern was evaluated on HeLa (cervical cancer) cell line using the MTT assay. The cells were incubated at 37°C for 72 hours in 5% CO_2_ humidified chamber in varying concentrations of FBS (0-10%) enriched traditional (DMEM) cell culture medium and ClearX9™. Cells grown with 0% and 10% FBS containing traditional (DMEM) cell culture medium were used as negative and standard controls (Fig. 3A, Fig. 3B).

**Fig 3.**
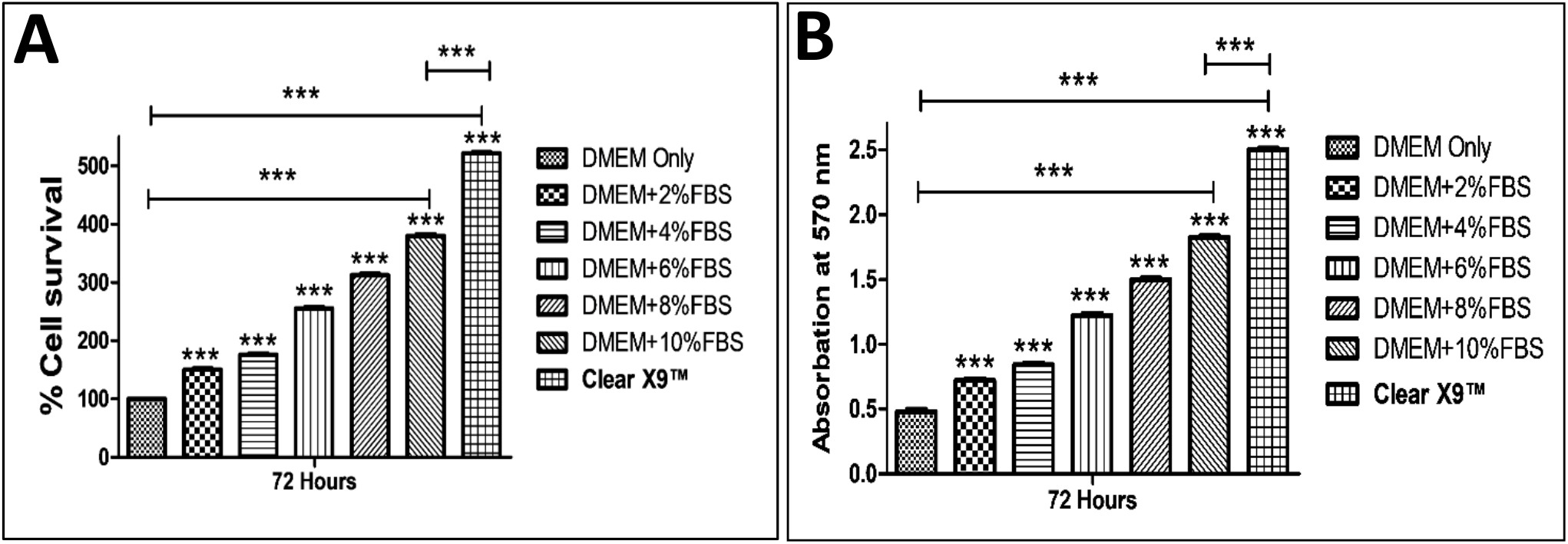
ClearX9™ enhanced cells proliferation rate of HeLa cells at 72 hours. **A)** Percentage of cell survival. **B)** Absorption (O.D. Value) at 570 nm. MTT cell Viability Assay was also performed on the HeLa cells. Briefly, cells (5 ×10^4^ cells/ml) were plated in 96 well plates and grown in presence of 0%FBS (Only DMEM), 2%, 4%, 6%, 8%, and 10% Fetal Bovine Serum (FBS) containing traditional complete medium and ClearX9™ for 72 hours. At the time of harvesting, MTT dye was added, and cells were incubated for 5 hours in a 37°C and 5% CO2 humidified chamber. The crystals formed in the process were dissolved in DMSO and OD was taken at 570nM. The data shown are an average of three parallel experiments, with each experiment performed in triplicate. ***p<0.001.

Results suggest that ClearX9™ significantly increased the proliferation rate of HeLa cells as compared to the 0-10% FBS containing traditional (DMEM) medium cells (Fig. 3A).

HeLa cells were plated at 7 × 10^5^ cells/ml in a 35mm^2^ culture dish and grown in the presence of 0-10% FBS containing traditional (DMEM) medium or ClearX9™ at 37°C in a 5% CO_2_ humidified chamber. HeLa cell morphology was visualized after 72 hours using a bright field microscope and all Images was captured by Zeiss Primovert inverted microscope at 20X magnification (Fig. 4).

**Figure 4.**
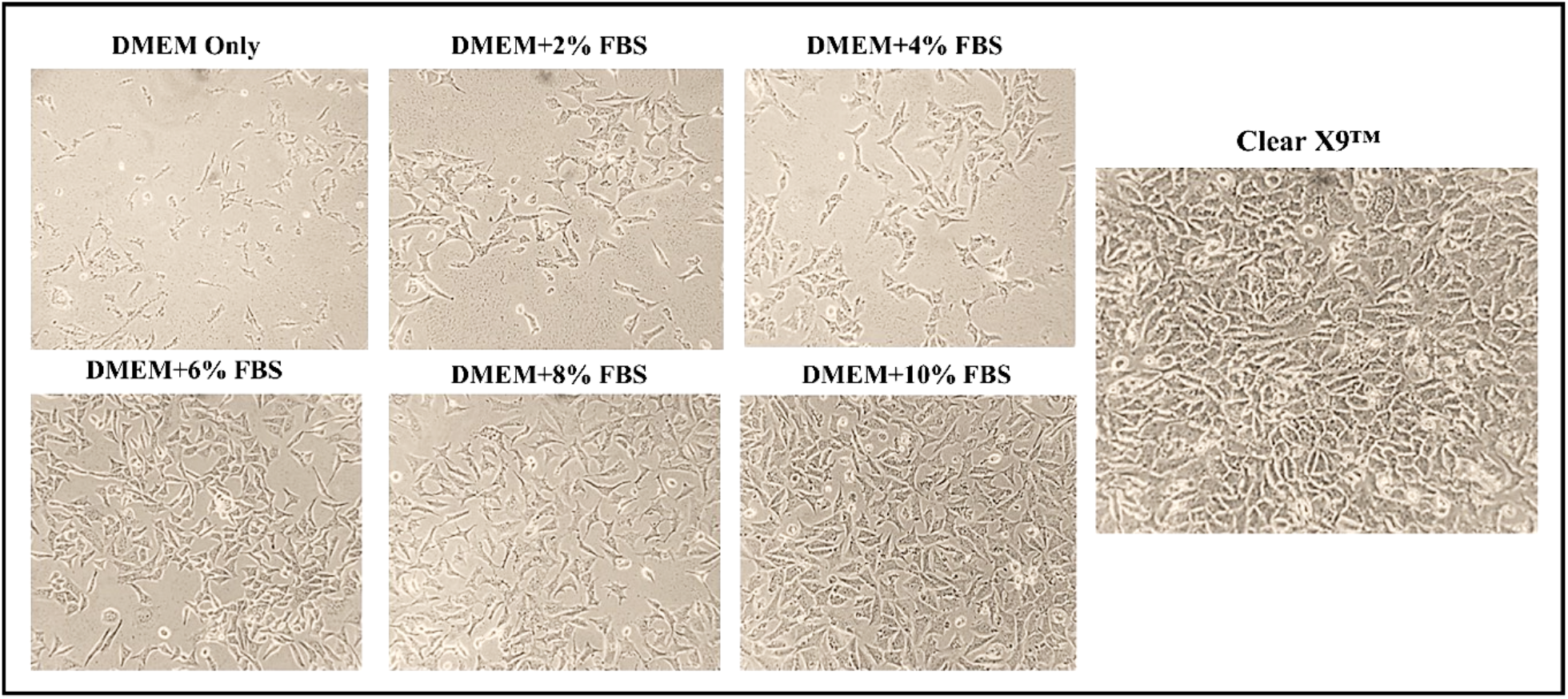
ClearX9™ induced growth rate of HeLa cells at 72 hours. 7 × 105 cells/ml cells were plated in a 35mm^2^ culture dish and grown in the presence of 0%FBS (Only DMEM), 2%, 4%, 6%, 8%, and 10% Fetal Bovine Serum (FBS) containing traditional complete medium and ClearX9™ and Cell Morphology was visualized after 72 hours using bright field microscope (Images were captured by Zeiss Primovert inverted microscope at 20X magnification).

The HeLa cell morphology data provides a clear evidence of maximum cell growth rate in ClearX9™ compared to varying concentrations of FBS. ClearX9™ significantly increased the growth rate of HeLa cells showing a healthy cell morphology compared to 0-10% of FBS-containing traditional (DMEM) medium (Fig. 4). HeLa cells were plated at 7 × 10^5^ cells/ml in a 35mm^2^ culture dish and grown in the presence of 0 and 10% FBS containing traditional (DMEM) medium or ClearX9™ at 37°C in a 5% CO_2_ humidified chamber. Cells grown with 0% and 10% FBS containing traditional (DMEM) cell culture medium served as a negative control and control in this experiment (Fig.5). HeLa cell morphology was visualized after 24, 48, and 72 hours using a bright field microscope, and all Images were captured by Zeiss Primovert inverted microscope at 20X magnification (Fig. 5).

**Fig. 5.**
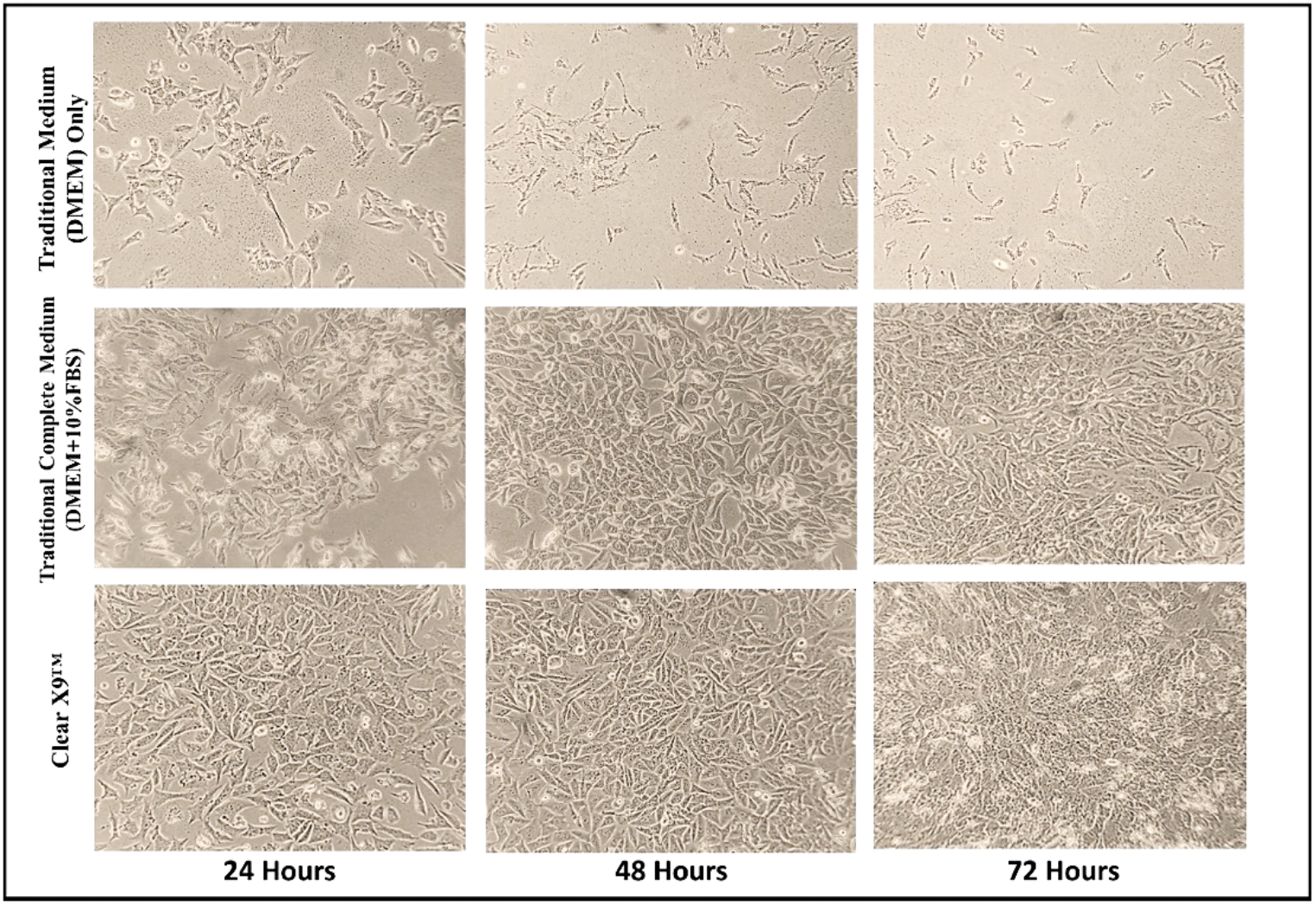
ClearX9™ increased the HeLa cell growth rate. 7 × 10^5^ cells/ml cells were plated in a 35mm^2^ culture dish and grown in the presence of 0%FBS (Only DMEM), complete traditional medium (DMEM+ 10%FBS), ClearX9™ and Cell Morphology was visualized after 24, 48, and 72 hours using bright field microscope (Images were captured by Zeiss Primovert inverted microscope at 20X magnification).

Results suggest that the maximum cell growth rate was observed in ClearX9™ and the minimum cell growth rate was observed in 0%FBS (Only DMEM) condition. ClearX9™ significantly increased the growth rate of HeLa cells with increasing time intervals with healthier cell morphology as compared to the 0% FBS (Only DMEM) and 10% FBS containing traditional (DMEM) medium cells (Fig. 5). These results are further supported by cell viability assay (Figs. 1 and 3).

Cell viability was evaluated on HeLa (cervical cancer cells), HEK293T (embryonic kidney transform cells), and Nthy Ori-3-1 (primary thyroid follicular transform epithelial cells) using MTT assay. The cells were incubated in the presence of 10% FBS containing traditional (DMEM/RPMI 1640) cell culture medium or ClearX9™ for 72 hours and grown at 37°C in a 5% CO_2_ humidified chamber. Hela and HEK 293T cells grown with 10% FBS containing traditional (DMEM) cell culture medium whereas Nthy Ori-3-1 cells grown with 10%FBS containing conditional (RPMI 1640) medium served as a control in this experiment (Fig. 6)

**Fig. 6.**
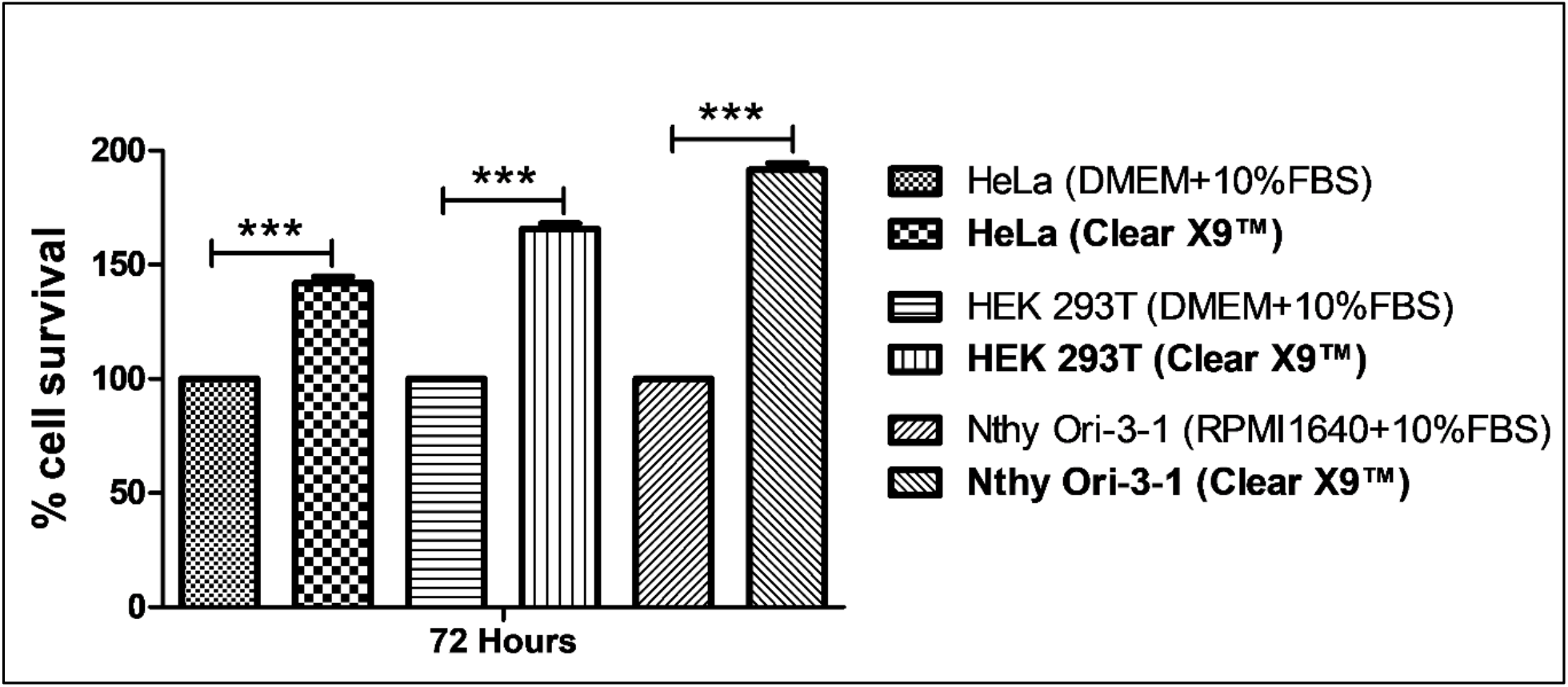
ClearX9™ enhanced cells proliferation rate of HeLa, HEK 293T, and Nthy Ori-3-1 cells at 72 hours. Cell Viability Assay was done by MTT on HeLa, HEK 293T, and Nthy Ori-3-1 cells. Briefly, cells (5 x104 cells/ml) were plated in 96 well plates and grown with 10% Fetal Bovine Serum (FBS) containing conditional (DMEM/RPMI 1640) complete medium and ClearX9™ for 72 hours. After the end of the stipulated time interval, MTT dye was added and incubated for 5 hours in a 37°C and 5% CO_2_ humidified chamber. The formed crystals were dissolved in DMSO and OD was taken at 570nM. The data shown are the mean from three parallel experiments, and each experiment was done in triplicate. ***p<0.001.

All cells were plated at 7 × 10^5^ cells/ml in a 35mm^2^ culture dish and grown in the presence of 10% FBS containing traditional complete medium or ClearX9™ at 37°C in a 5% CO_2_ humidified chamber. Hela and HEK 293T cells were grown with 10% FBS containing traditional (DMEM) cell culture medium whereas Nthy Ori-3-1 cells grown with 10%FBS containing conditional (RPMI 1640) medium served as a control in this experiment (Figs. 6 & 7). All cell morphology was visualized after 72 hours using a bright field microscope and all Images were captured by Zeiss Primovert inverted microscope at 20X magnification (Fig. 7). Results suggest maximal cell growth rate in ClearX9™ compared to the cells grown in 10%FBS (Fig. 6). ClearX9™ increased the growth rate of cells significantly with healthy cells showing up in the culture indicating stress-free growth compared to the 10% FBS enriched traditional (DMEM/RPMI-1640) medium cells (Fig.7 and Fig.8).

**Fig. 7.**
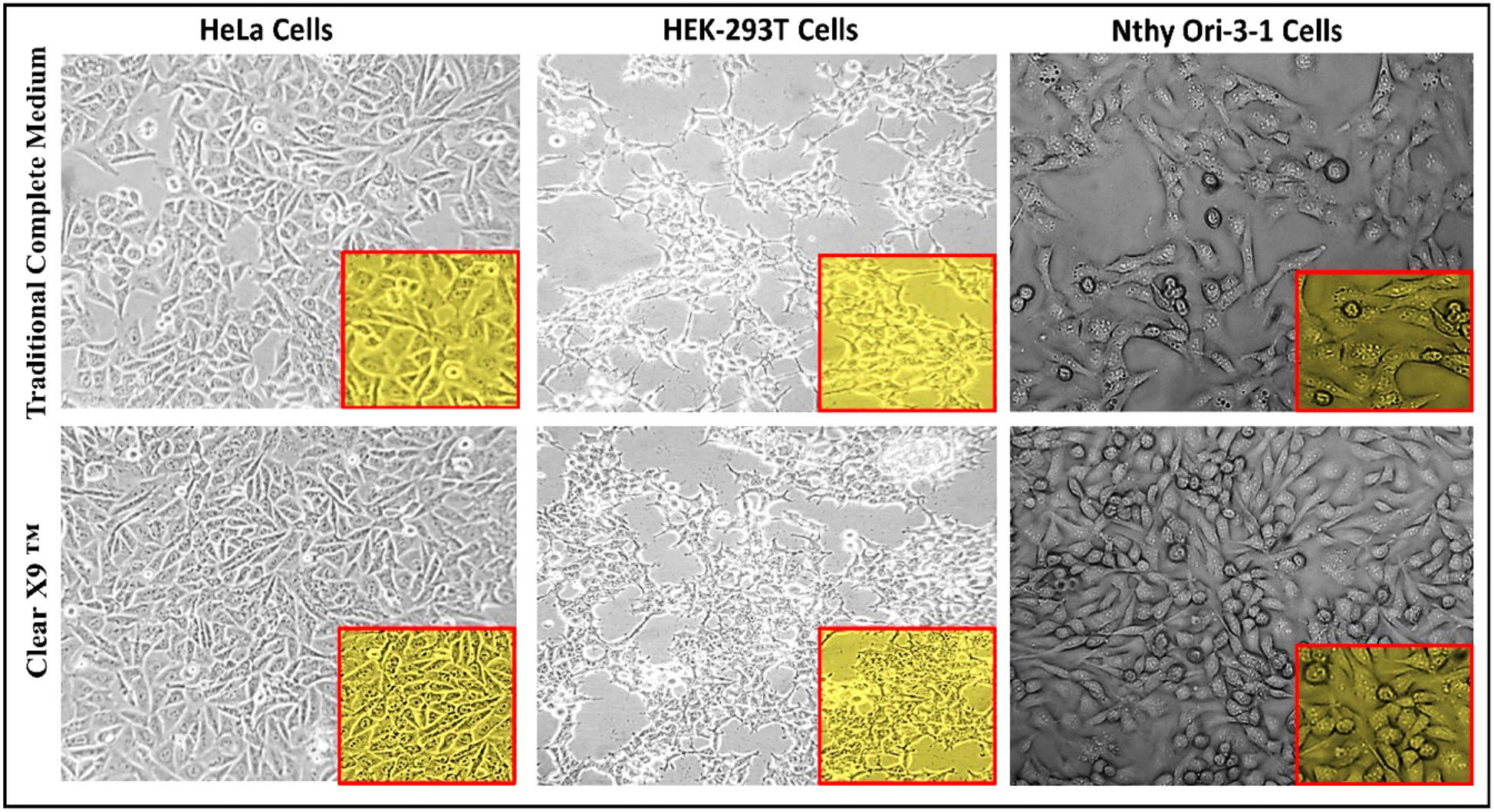
ClearX9™ increased the HeLa, HEK-293T, and Nthy Ori-3-1 cells growth rate at 72 hours. 7 × 10^5^ cells/ml cells were plated in a 35mm^2^ culture dish and grown in the presence of a complete traditional medium (DMEM/RPMI 1640+10%FBS) and ClearX9™ and cell morphology was visualized after 72 hours using a bright field microscope (Images were captured by Zeiss Primovert inverted microscope at 20X magnification

**Fig 8.**
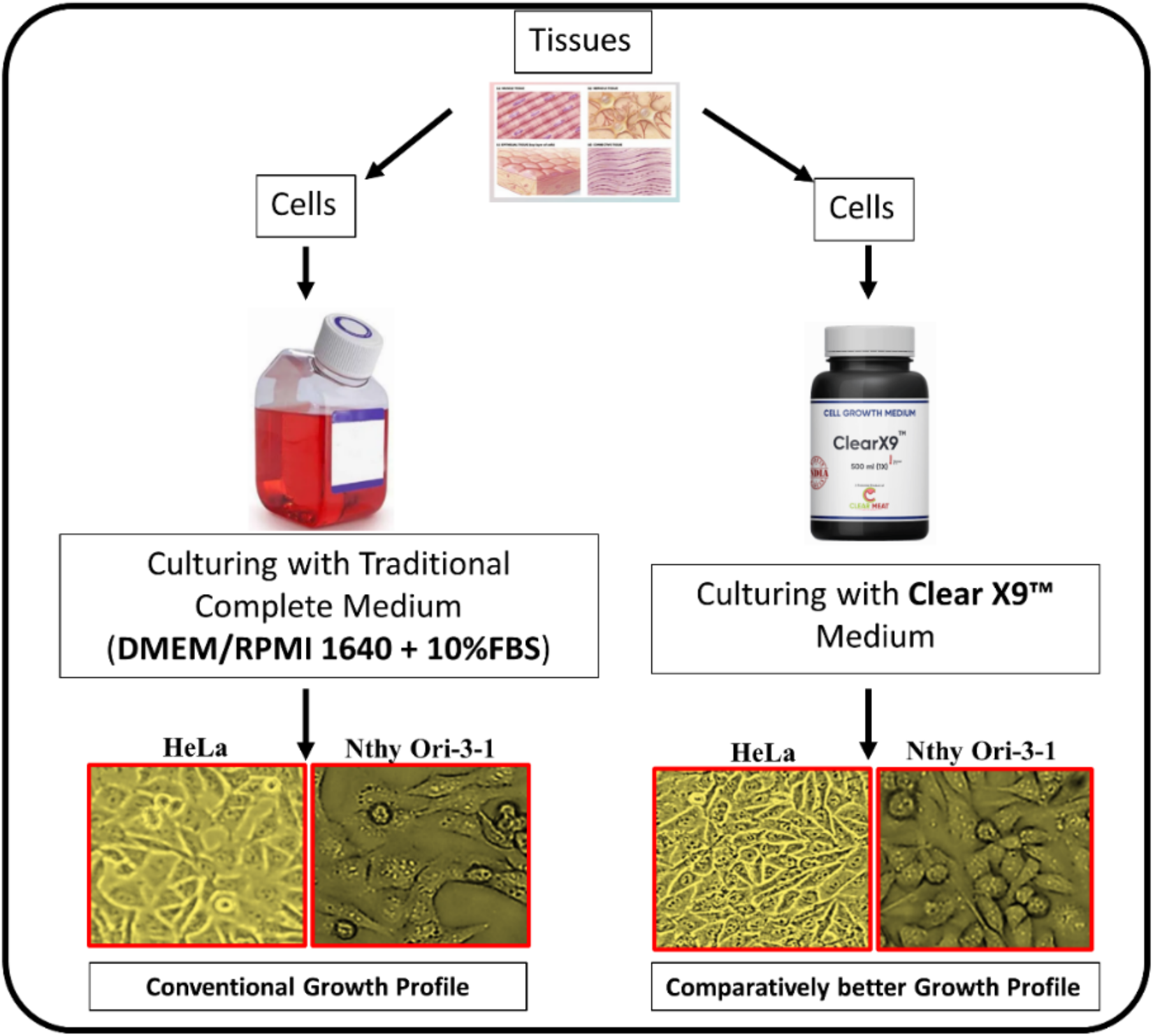
Mammalian cancer cells grown in ClearX9™ & 10% FBS enriched cell culture medium. The conventional cell growth is clearly distinguished from the ClearX9™ supported cell growth

## 4. DISCUSSION

The origin of FBS can be traced back to the practice of collecting blood from pregnant cows during slaughter, which was found to contain high levels of growth factors and other components like amino acids, sugars, lipids, hormones, and low gamma globulin that support cell growth. FBS is the plasma left after the natural coagulation of the blood and contains low levels of antibodies compared to other animal sera. Due to fluctuation in the procurement and distribution pipeline, the price of FBS has been volatile with a change in international trading policies and changes in the beef processing industry increases the cost of fetal bovine serum in certain countries.

As animal are not raised exclusively for the serum extraction, the meat supply and demand changes may change the cost of the fetal bovine serum. Even if FBS is a popular essential component of cell culture media, particularly adherent cells, the solution is far from perfect, as it includes physical torture to animals, variability between batches, high cost and the risk of transmitting pathogens to humans.

The disadvantages can also arise from ill-defined description of components, presence of adverse factors such as endotoxin, mycoplasma, viral contaminants, or prion proteins (Gstraunthaler et al., 2013).

With an estimated millions of pregnant cows slaughtered for serum, loose regulation of FBS market resulting in a dilution of the FBS quality, cruelty to animals and huge methane emissions from cattle industry, (Sinke et al., 2023; Treich, 2021; Tuomisto and Teixeira de Mattos, 2010; Tuomisto and Teixeira De Mattos, 2011), there is a pressing need for innovative solutions.

Researchers have been exploring possibilities in the form of serum-free or serum-reduced media supplemented with growth factors, cytokines, or other additives, and plant based or synthetic serum supplements to provide consistent and reproducible cell culture conditions while minimizing animal slaughter. However, efforts seem to be largely localized and lab based with little impact on the global FBS market.

In this context, the development of ClearX9™ assumes significance, as ClearX9™ meets performance indicators of cell growth parameters in comparison to the cells grown in the FBS enriched media. Cells cultured in ClearX9™ shows clarity of cytoplasm indicating stress-free environment compared to traditional complete medium condition.

ClearX9™ has been designed to provide a ready-to-use, natural component-based product, with adequate supply of amino acids, vitamins, lipids, plant pigments, and natural growth promoters for growth of animal cells in-vitro.

In this report, we provide the first experimental evidence of using ClearX9™ to grow HeLa (cervical cancer cells), HEK293T (embryonic kidney transform cells), and Nthy Ori-3-1 (primary thyroid follicular transform epithelial cells).

Results indicate that ClearX9™ mediated proliferation of cells retains a healthy growth and a healthy morphology, compared to the cells grown in FBS enriched cell culture medium. Data strongly suggest ClearX9™ as a complete and sustainable cell culture medium for biotechnology and lab grown meat industries. The cellular morphology metric, based on vacuole concentration, has been traditionally used as a standard parameter for defining health of a cell. Microscopic studies suggest a vacuole free cytoplasmic presentation in cells growing in ClearX9™. A less clear cytoplasmic presentation was found in cells grown in 10% FBS enriched medium thereby indicating presence of several dead cells (Crowley et al., 2016; Thong et al., 2016).

In Nthy Ori-3-1 cells, vacuoles were found to be present in 10%FBS enriched RPMI 1640 complete medium indicating stressful condition within cells in comparison to the stress-free cytoplasmic environment observed in ClearX9™ grown cells.

This study suggests potential of ClearX9 as a credible FBS alternative in the biotech and cell cultivated meat sectors. More work is required to study the impact of ClearX9™ in sectors like vaccine manufacturing, regenerative medicine, biomanufacturing, and toxicology. Furthermore, there is a need to perform in depth pathway analysis to answer cell growth and cell health questions at the molecular level. In future, it would be interesting to know how genes are up/down regulated in a range of FBS vs. ClearX9™ combinations.

## ACKNOWLEDGEMENT

Clear Meat Pvt. Ltd. expresses sincere gratitude to the key investors Gastrope Accelerator Group (supported by Miseltoe), BRINC Group, and Artesian HB Group, and Gene Kim for funding this work. We warmly thank all scientists and students who tested the ClearX9™ culture medium and shared their valuable feedback.

## AUTHOR’S CONTRIBUTIONS

The concept and formulation of ClearX9™ medium was conceived by PKD and supervised by SM for optimization. All the experiments were jointly performed by SG and NV. The manuscript was written by SG and PKD. All authors approved data presented in this paper.

SG: Sumit Gautam
NV: Neeraj Verma
SM: Siddharth Manvati
PKD: Pawan K. Dhar

